# Light-guided molecular patterning for programmable multiplexed single-molecule manipulation

**DOI:** 10.1101/2025.04.30.651527

**Authors:** Hansol Choi, Andrew Ward, Wesley P. Wong

## Abstract

Single-molecule force spectroscopy enables the detailed probing of molecular interactions, providing new insights into molecular mechanisms—yet studying biology “one molecule at a time” can lead to throughput challenges that limit applications. While multiplexed single-molecule assays can address these issues, suitable functionalization of surfaces is required, which remains a technical challenge—many commonly used approaches are constrained by random and sparse biomolecules arrangements, limiting programmability and throughput. An ideal anchoring method would enable (i) high surface densities to maximize throughput, (ii) precise control of spatial position and molecular identity for maximum control, (iii) covalent linking for high force application, and (iv) efficient patterning without the need for expensive facilities to maximize accessibility. To achieve these aims, we have developed a light-guided surface patterning method that can covalently organize oligonucleotides (oligos) without the need for lithographic equipment. Oligos with 3-Cyanovinylcarbazole (CNVK) nucleoside are crosslinked by UV patterns reflected through a digital micromirror device (DMD), with beads arranged accordingly. To demonstrate compatibility with established single-molecule methods, we performed single-molecule force spectroscopy experiments on patterned coverslips, using both magnetic tweezers and hydrodynamic-based systems. Our light-guided approach provides a scalable and accessible solution for biomolecular patterning that allows precise control over molecular identity and spatial positioning, enabling high-throughput measurements in single-molecule research.

## Introduction

Single-molecule force spectroscopy has become an essential tool in biophysics for analyzing force driven biological processes^1–4^ including receptor binding^5,6^ and motor protein activity^7^. While these approaches are leading to new biological insights, studying biology “one molecule at a time” has throughput challenges that can limit applications. To address this, multiplexed single-molecule approaches have been developed, including parallel magnetic tweezers^8,9^, centrifuge force microscopy^10–12^, acoustic force spectroscopy^13^, and hydrodynamic force spectroscopy^14–16^. These methods apply precise forces to multiple molecules simultaneously, enhancing throughput by tracking multiple beads tethered to biomolecules on surfaces. Such methods can expand the applications of single-molecule analysis, facilitating the analysis of complex samples, library screening, and characterization of molecular heterogeneity. The throughput and programmability of these multiplexed methods largely depend on the characteristics of surface functionalization^17^—the spatial arrangement and density of biomolecules and beads on the surface, as well as the specificity of linkage chemistries. Yet common functionalization approaches are often constrained by either random and sparse distributions of beads, limiting throughput, or the need for expensive lithographic equipment, reducing accessibility. Furthermore, many methods rely on non-covalent interactions (e.g., adsorption), limiting the range of forces that can be applied during experiments^18^.

Here, we present a method that can precisely organize biomolecules at single-molecule resolution without the need for photomasks or lithographic instruments, enabling spatially resolved single-molecule experiments on a patterned surface. Oligos are patterned on a flow cell by employing 3-Cyanovinylcarbazole (CNVK) that can be covalently crosslinked upon exposure to UV light (365 nm)^19,20^, with illumination of patterned UV achieved through the use of a digital micromirror device (DMD)^21,22^. To demonstrate programmability and spatial control, we create diverse patterns of biomolecules such as square, and hexagonal lattice pattern with various spacings. Moreover, oligos with different sequences can be sequentially patterned on the flow cell, enabling different molecular species to be precisely arranged for spatially resolved experiments. To demonstrate how this patterning approach can enable highly parallel single-molecule force spectroscopy measurements, we use magnetic tweezers and flow-based force spectroscopy to stretch and unzip DNA constructs on multiplexed patterned surfaces.

## Results

### Principles of light-guided molecular patterning

Our light-guided molecular patterning method enables the specific arrangement of oligos onto a solid surface; these oligos can furthermore be used to tether beads to targeted position for single-molecule force studies (Fig. 1a). Oligos are covalently attached to the surface via light-activated CNVK crosslinkers, with their spatial arrangement determined by the illuminated UV pattern (Fig. 1b). To passivate the surface and prepare it for patterning, 17 nucleotide (nt) base oligos modified with DBCO-PEG13 are covalently conjugated to the azide-functionalized coverslip using copper-free click chemistry. Next, patterning oligos are added that can transiently hybridize to the base oligos (Supplementary Fig. 2). The patterning oligos consists of a 14 nt region, which includes a CNVK nucleoside, for hybridization to the base oligos, and a 55 nt functional region designed to capture DNA constructs for tethering beads. Upon UV illumination, the CNVK nucleosides crosslink to the thymine bases in the base oligos within seconds, enabling covalent oligos patterning. Transiently hybridized oligos that are not exposed to UV are denatured and washed with formamide to ensure oligo patterns match the UV pattern.

**Figure 1.**
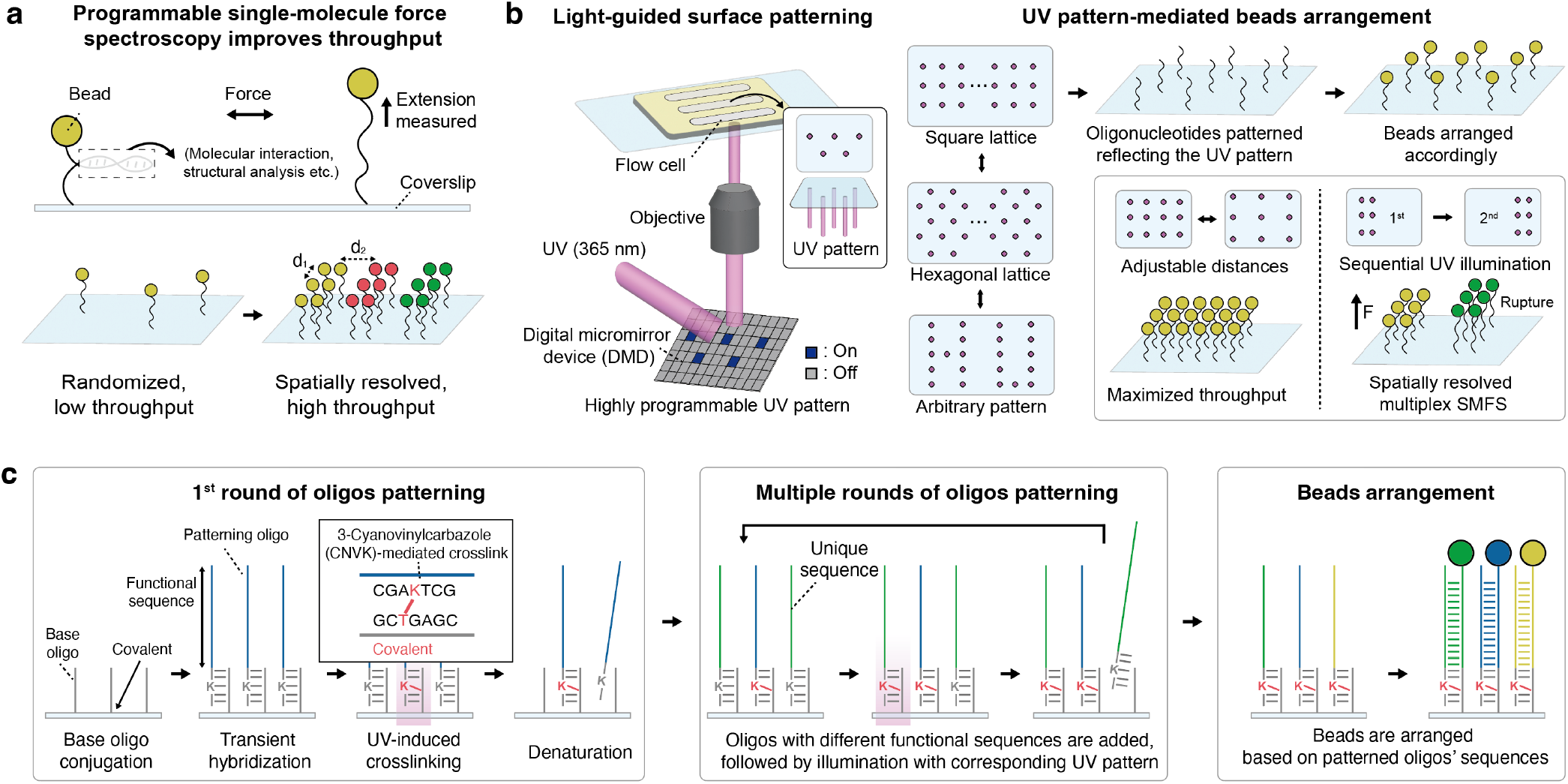
Spatially resolved single-molecule force spectroscopy achieved by light-guided surface patterning. (a) The position of beads is precisely controlled to minimize interactions between adjacent beads and to enable multiplexed single-molecule force spectroscopy. (b) UV light is patterned through a digital micromirror device (DMD) to crosslink oligos where UV is illuminated. The distance between adjacent beads is programmable, allowing distinct oligos to be spatially resolved for multiplex experiments. (c) Base oligos are covalently conjugated to the surface, followed by hybridization with patterning oligos containing CNVK. Where UV is illuminated, patterning oligos and base oligos are crosslinked, and oligos that are not exposed to UV are denatured. This patterning process can be repeated to conjugate patterning oligos with different functional sequences. Beads are arranged based on the functional sequences of the crosslinked oligos.

Regarding the patterning resolution, the nominal size of the illumination spot, not including diffraction, is approximately 390 nm by 390 nm when the UV is illuminated through a 20X objective with a single micromirror “pixel” activated. To experimentally characterize the spot size, taking into account the point-spread function of our imaging/illumination system, fluorescent oligos complementary to the functional region were hybridized, and the fluorescence intensity of the resulting pattern was analyzed. The observed intensity was fitted to a Gaussian, resulting in a full width at half maximum (FWHM) of 992 nm (Supplementary Fig. 3).

To achieve sequence programmability, the flow cell can be patterned with multiple patterning oligos with unique functional sequences by repeating the cross-linking and washing procedure multiple times, once for each sequence. This could be used to enable, for example, multiplexed spatially resolved single-molecule force spectroscopy. The functional sequences of the patterning oligos were analyzed with NUPACK^23^ to generate orthogonal sequences and minimize secondary structure.

To tether beads to the patterned oligos, DNA constructs that include a 55 nt region that is reverse complementary to these functional sequences are conjugated to streptavidin-coated beads, which are then incubated with the surface (Fig. 1c). Alternatively, constructs with biotin modifications can be hybridized to patterning oligos, followed by streptavidin-coated bead incubation in the flow cell (Supplementary Fig. 4).

### Fabrication of programmable bead arrays

When performing single-molecule force spectroscopy, the optimal distance between adjacent beads is determined by the lengths of the constructs connecting beads to the surface and the size of beads; this is particularly important with magnetic tweezers to prevent interactions between tethered beads^18^. While previous methods often require the creation of a photomask and/or the use of lithographic methods, which can be time consuming, our light-based approach enables arbitrary patterns to be generated “on the fly”.

To demonstrate this flexibility and precise spatial positioning, beads arrays were fabricated in both hexagonal and square lattice patterns with three different bead spacings (Fig. 2a). The primitive vector lengths or minimal distances between adjacent beads were set at 11.5 µm, 15.3 µm, and 19.2 µm for both square and hexagonal lattice patterns. The size of the illumination patch for each pattern was set to a nominal 780 nm by 780 nm square (2x2 pixels), sufficient to tether a single bead with a diameter of 2.8 µm on each patch and to prevent other beads from tethering due to steric hinderance. Streptavidin beads were conjugated with 55 bp biotin-modified oligos that were reverse complementary to the functional sequence of patterning oligos on the flow cell surface.

**Figure 2.**
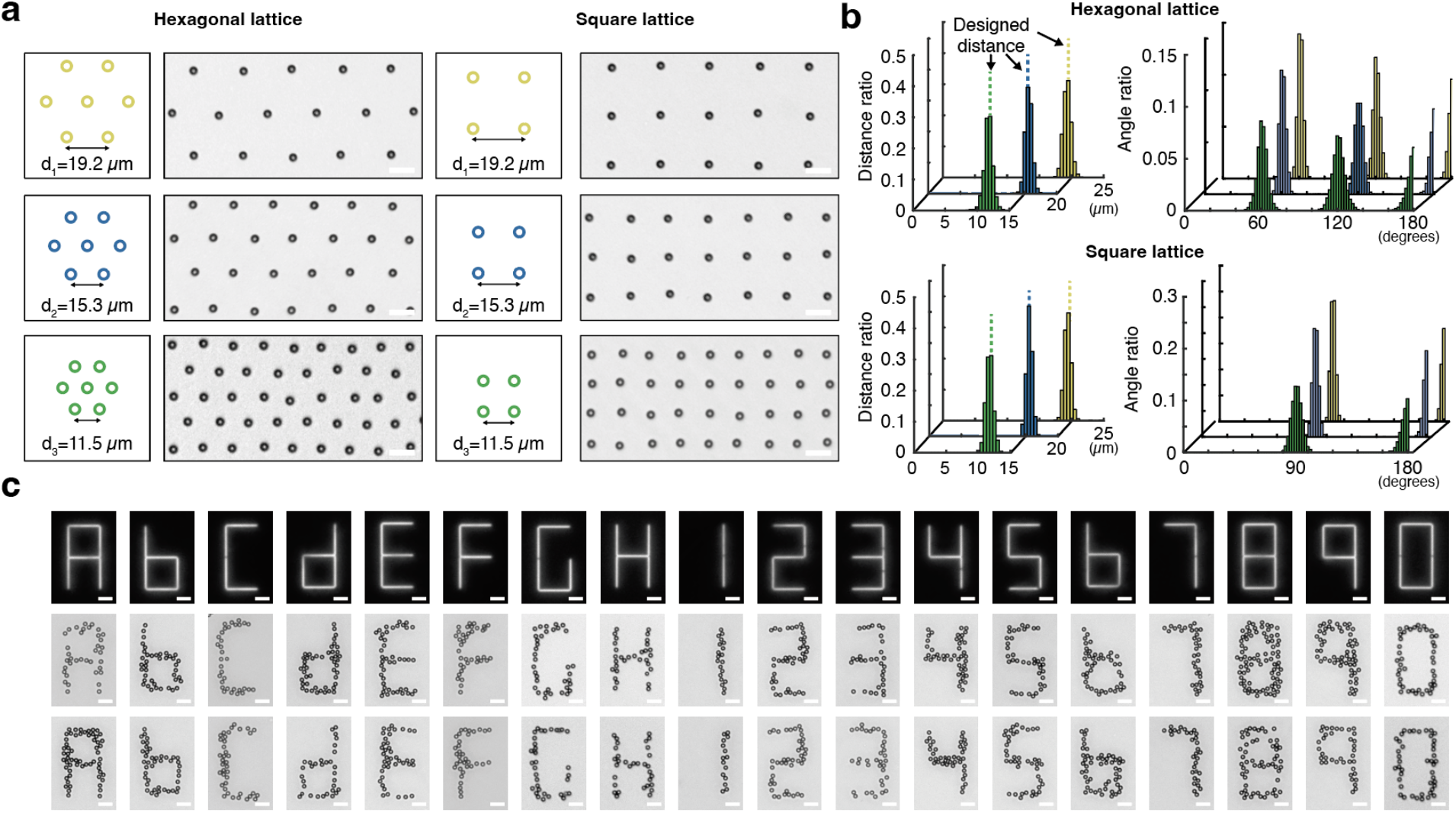
Diverse bead patterns enabled by a highly programmable patterning method. (a) Beads are organized in hexagonal and square lattice pattern with minimal distances of 11.5 µm, 15.3 µm, and 19.2 µm. (b) Probability density of distance distribution and angle distribution are all patterns are analyzed. (c) Various beads patterns in 7 segments are fabricated. Scale bar, 10 µm.

Subsequently, the oligo-conjugated beads were introduced into the flow cell using a syringe pump with repeated cycles of infusion and withdrawal to increase the probability of tethering a bead to each spot. Untethered beads were washed away with buffer, and the remaining patterned beads were analyzed. Beads positions were measured using an edge detection algorithm employing the Sobel approximation^24^ and three fields of view of the DMD were employed for the analysis of each condition (Supplementary Fig. 5-10). The X and Y coordinates of the bead centers were measured to analyze distance and angle distributions (Supplementary Fig. 11). Although distances between all beads were calculated, a distance cut-off of 1.3 times the length of the primitive vector was applied to specifically examine distances distribution between adjacent beads (Fig. 2b). The mean and standard deviation (s.d.) of measured distances for the square bead lattices were 11.49 ± 0.58 µm, 15.31 ± 0.51 µm, and 19.19 ± 1.17 µm, in excellent agreement with the designed distances. Similarly, the hexagonal bead lattices exhibited distances of 11.5 ± 0.67 µm, 15.37 ± 0.62 µm, and 19.07 ± 0.59 µm (mean ± s.d.), also exhibiting good quantitative agreement. Angle distributions were analyzed by setting a reference bead as a vertex for each angle, with two additional adjacent beads employed to measure the angle between them. The same distance cut-off was applied when selecting the two adjacent beads, and angle measurements were made for every bead as a vertex. All angle measurements gave the expected values. Specifically, for the hexagonal bead lattice with 11.5 µm spacing, measured angles were 60 ± 4.1°, 120 ± 5.1°, and 175.8 ± 3.6° while those with 15.3 µm showed 59.9 ± 3.7°, 120 ± 3.6°, 177.1 ± 2.5°, and those with 19.2 µm spacing showed 60 ± 2.1°, 120 ± 2.7°, 177.7 ± 1.8° (mean ± s.d.) respectively. The angles measured from the square lattice bead arrays with a 11.5 µm spacing were 90 ± 4.1°, and 176 ± 3°. Similarly, angles measured from the square lattices with 15.3 µm and 19.2 µm spacings were 90 ± 2.5°, 177.7 ± 2.2°, and 89.6 ± 6°, 177.7 ± 3.6° (mean ± s.d.), respectively.

As a further demonstration of the high spatial programmability of our patterning technology, various alphabets and numbers in seven segments, where each line segment had a length of 22.6µm, were fabricated with beads (Fig. 2c). In addition to the demonstrated patterns, any arbitrary patterns can be fabricated without any photomasks since the illumination pattern generated by the DMD can be immediately adjusted in response to an input image file.

### Multiplexed patterning with oligos of different sequences

Our patterning method offers high molecular programmability, enabling the spatial organization of molecules with multiple distinct identities. By repeating the patterning process described above to create oligo patterns with distinct sequences, multiple different species of biomolecules or beads with different surface chemistries can be specifically and precisely attached to the surface (Fig. 3a). We demonstrated this capability by patterning two distinct molecular species onto a surface. First, patterning oligos with sequence 1 were arranged on the surface in a square lattice pattern by illuminating the desired spots with UV light as before. This was then followed by a washing step and a second round of patterning. Specifically, a second type of patterning oligos with sequence 2 was introduce into the flow cell; during incubation, another square lattice UV pattern, shifted to illuminate the center of the unit cell of the initial lattice, was used to crosslink the second patterning oligos; this process was then followed by a stringent washing step to remove all non-crosslinked oligos. Two different types of fluorescent beads were then tethered to these two distinct sequences. Beads were created by first incubating streptavidinated beads with biotinylated oligos that were reverse complementary to either functional sequence 1 or 2, followed by incubation with dyes with biotin or NHS-ester modifications; a different dye was used for each type of bead. Incubation of these beads with the surface resulted in efficient tethering of beads to the lattice-organized patterning oligos on the flow cell surface, with spatial organization based on sequence. After washing away the untethered beads, the resulting bead patterns were imaged with bright field and two distinct fluorescence channels to demonstrate the capability of our method to precisely position multiple distinct molecular targets (Fig. 3b, Supplementary Fig. 12).

**Figure 3.**
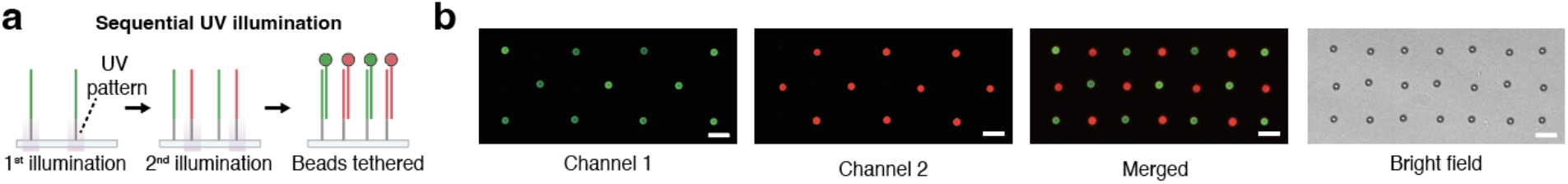
Spatially resolved multi-sequence bead tethering by multiple rounds of patterning. (a) Two unique UV patterns are employed to conjugate two distinct patterning oligos. (b) Beads covered with different dyes are attached to corresponding patterning oligos. Images taken with two different fluorescence channel and bright field are shown. Scale bar, 10 µm.

### Force spectroscopy with magnetic tweezers on patterned surfaces

Single-molecule force measurements with magnetic tweezers were conducted on the patterned flow cell. To facilitate repeated rupture measurements at single-molecule resolution magnetic beads were tethered to the surface by DNA nanoswitches^11,25–27^, 7.2 kb long DNA constructs with two biomolecules of interest attached at different positions along the length ^9,21,22^(Fig. 4a). This construct forms a loop when these biomolecules bind to each other, and switches to a linear state when the bond breaks, resulting in an easily observable increase in length. For our demonstration, nanoswitches with two unzipping oligos designed to hybridized to different positions on the DNA construct with overhangs that were reverse complementary were used to measure the force-driven DNA unzipping transition^28^. Four different types of DNA nanoswitches were created, with two different unzipping oligo sequences with GC contents of 48% and 31%, designed to measure different unzipping forces, and two different loop sizes of 1.1 µm and 0.65 µm, designed to produce different changes in length upon opening (Supplementary Fig. 13). The construct was prepared by linearizing circular single-stranded DNA, followed by hybridization of 120 60-bp backbone oligos to make the construct double-stranded. The fabrication efficiency and the fraction of looped constructs were analyzed with gel electrophoresis, reaching up to 97.5% (Supplementary Figure 14–15). The looping fraction varied depending on the unzipping oligo sequences and the loop size of the construct; unlooped constructs could result from synthetic errors in the unzipping oligos^29^ or from single-stranded overhangs hybridizing with other unzipping oligos in the solution.

**Figure 4.**
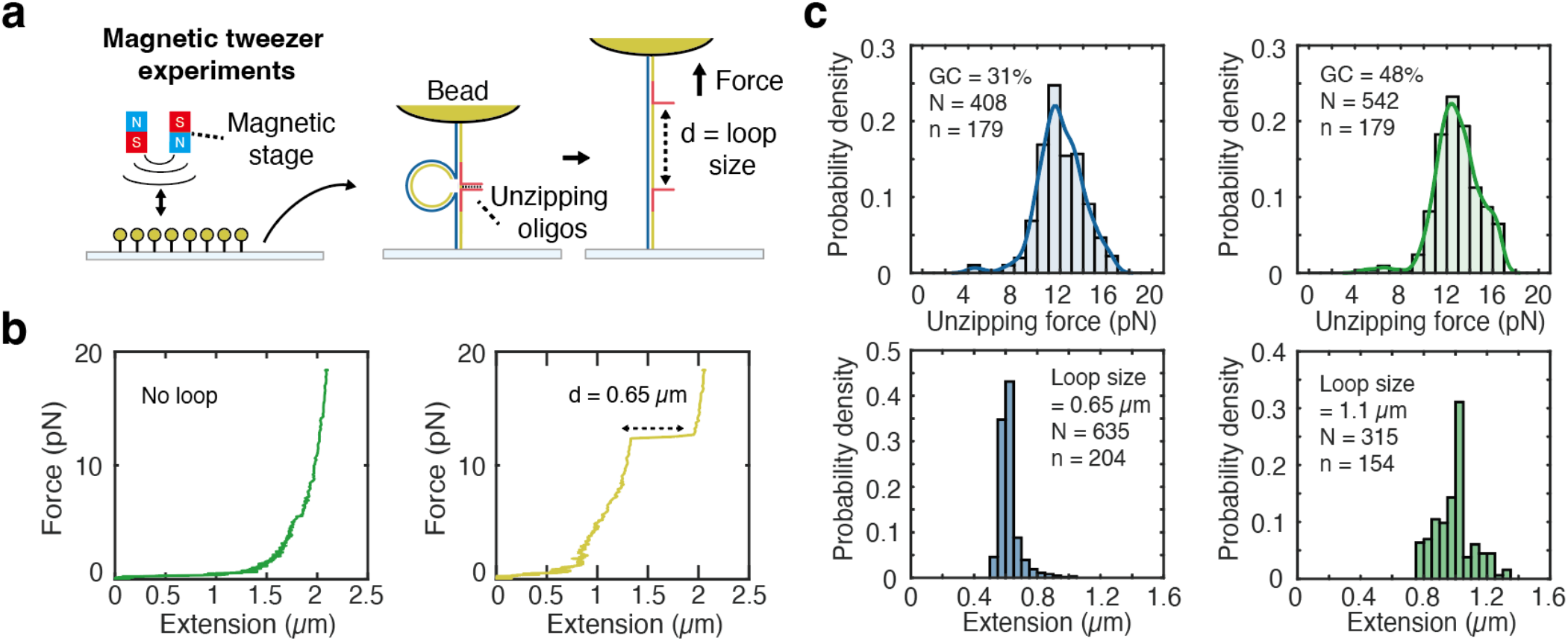
Magnetic tweezers experiments conducted on patterned coverslips. (a) Magnetic force was applied by lowering magnets to the sample on the patterned coverslip. DNA nanoswitch constructs with loop sizes of 1.1 µm and 0.65 µm, and unzipping oligos with different GC contents were designed. (b) Force-extension curves of the DNA nanoswitch constructs. The left panel shows the construct without unzipping oligos, and the right panel show the construct with a loop size of 0.65 µm and 48% GC content. (c) Histograms of unzipping force with GC content of 31% (top left), and 48% (top right) are shown. Histograms of extension of DNA nanoswitch due to unzipping with loop sizes of 0.65 µm (bottom left) and 1.1 µm (bottom right) are shown. N represents the total number of measurements colslected across n molecules.

For single molecule experiments, each spot would ideally be occupied by a single construct to decrease the chance of forming multiple tethers. To help facilitate this, a nominal illumination spot size of 390 nm was used, the smallest patch size achievable on our system with the 20X objective. Nanoswitch constructs with hybridization regions reverse complementary to the patterning oligos on one end and biotin-modified oligos at the other end for coupling to magnetic beads were incubated in the flow cell. Given that the number of crosslinked patterning oligos is determined by UV exposure^30^, we further optimized the number of single tethers on the patterned glass coverslip by controlling the UV power (Supplementary Fig. 16).

The force on the molecular tethers was calibrated using the equipartition theorem^31–33^ by measuring the mean square displacement of the beads at multiple different magnet heights (Supplementary Note 1). At the start of each measurement cycle the DNA nanoswitch constructs were putatively looped and under low magnetic force; the force was then gradually increased up to 18.4 pN to unzip the construct and open the loop. Loop opening was marked by an abrupt change in extension that provided a clear molecular signature to characterize and verify the unzipping transition (Fig. 4b). To filter out multiply tethered beads from those tethered with a single nanoswitch construct, we selected beads with tether lengths of at least 1.8 µm, and with loop sizes of at least 70% of the designed distances (1.1 µm or 0.65 µm) for analysis, based on measurements obtained using magnetic tweezers. DNA constructs with two different loop sizes were used, and a square lattice pattern with a minimal distance of 15.3 µm was used to prevent interaction between adjacent superparamagnetic beads during the application of magnetic force. The mean and standard deviation of unzipping forces for constructs with 48 % GC content and loop sizes of 1.1 µm and 0.65 µm were 12.6 ± 2.2 pN and 13.1 ± 1.6 pN respectively with a mean loading rate of 30 ± 7 pN/s (mean ± s.d.), showing unzipping forces are consistent regardless of the loop size. The unzipping forces for constructs with 31% GC content and loop sizes of 1.1 µm and 0.65 µm were 12.1 ± 2.1 pN and 12.0 ± 1.9 pN each with a mean loading rate of 27.1 ± 8.3 pN/s (mean ± s.d.). When the unzipping forces were calculated considering only the unzipping sequences, constructs with 31% GC content had a mean unzipping force of 12.0 ± 2.0 pN, and those with 48% GC content had a mean unzipping force of 12.9 ± 2.0 pN (mean ± s.d.). The ratio of those forces matched previously reported research using the same sequences^11^. Additionally, the extended lengths of DNA nanoswitches due to unzipping were 992 ± 120 nm for the constructs with a loop size of 1.1 µm, and 620 ± 65 nm (mean ± s.d.) for those with a loop size of 0.65 µm. The observed measurements were in good agreement with the expected values for dsDNA stretching under forces of 12 to 13 pN.

### Hydrodynamic force spectroscopy on patterned surfaces

To demonstrate that the method can be applied to various force spectroscopy experiments, single-molecule force spectroscopy with hydrodynamic force was performed on the patterned flow cell (Fig. 5a). The flow cell was connected to a syringe pump through polyetheretherketone (peek) tubing to apply hydrodynamic force based on the flow speed. The force calibration was conducted by calculating the drag force on the beads and the system’s geometry, both of which were used to determine the tension applied to the construct^35^ (Supplementary Note 1, Supplementary Fig. 1). The DNA nanoswitches used in the magnetic tweezer experiments were employed for hydrodynamic force spectroscopy, with beads tethered to each spot following the same protocol (Supplementary Fig. 17). To distinguish singly tethered beads from those with multiple tethers, we applied the same filtering criteria as in the magnetic tweezer experiments, determining tether lengths by comparing bead positions during flow infusion and withdrawal. With a loading rate of 1.89 pN/s for DNA nanoswitches with a loop size of 0.65 µm, the unzipping forces of the unzipping oligos with 31% GC content were 10.35 ± 1.27 pN, while those with 48% GC content were 11.07 ± 1.23 pN (mean ± s.d.; Fig. 5b). For the constructs with a loop size of 1.1 µm, the loading rate was 1.99 pN/s, and the unzipping forces of the unzipping oligos with 31% GC content were 10.4 ± 1.41 pN while those with 48% GC content were 11.44 ± 1.32 pN (mean ± s.d.; Supplementary Fig. 18). The ratio between the two unzipping forces were in good agreement regardless of the loop sizes. When the unzipping forces were calculated regardless of loop sizes, the ratio of unzipping forces between constructs with 31% GC content, measured using magnetic tweezers and flow experiments, was 1.16, while that for constructs with 48% GC content was 1.15, which is consistent with principles of dynamic force spectroscopy as lower loading rates generally decrease the value of the most probable rupture force^35–37^. Also, the nanoswitch length extensions along the X axis due to unzipping events were measured as 1060 ± 91 nm and 1102 ± 194 nm (mean ± s.d.) for constructs with a loop size of 1.1 µm and GC content of 48% and 31%, respectively. Similarly, extensions of 639 ± 127 nm and 635 ± 107 nm (mean ± s.d.) were observed for constructs with a loop size of 0.65 µm and GC content of 48% and 31%, showing that the extensions are consistent regardless of the unzipping oligos’ sequences (Fig. 5b).

**Figure 5.**
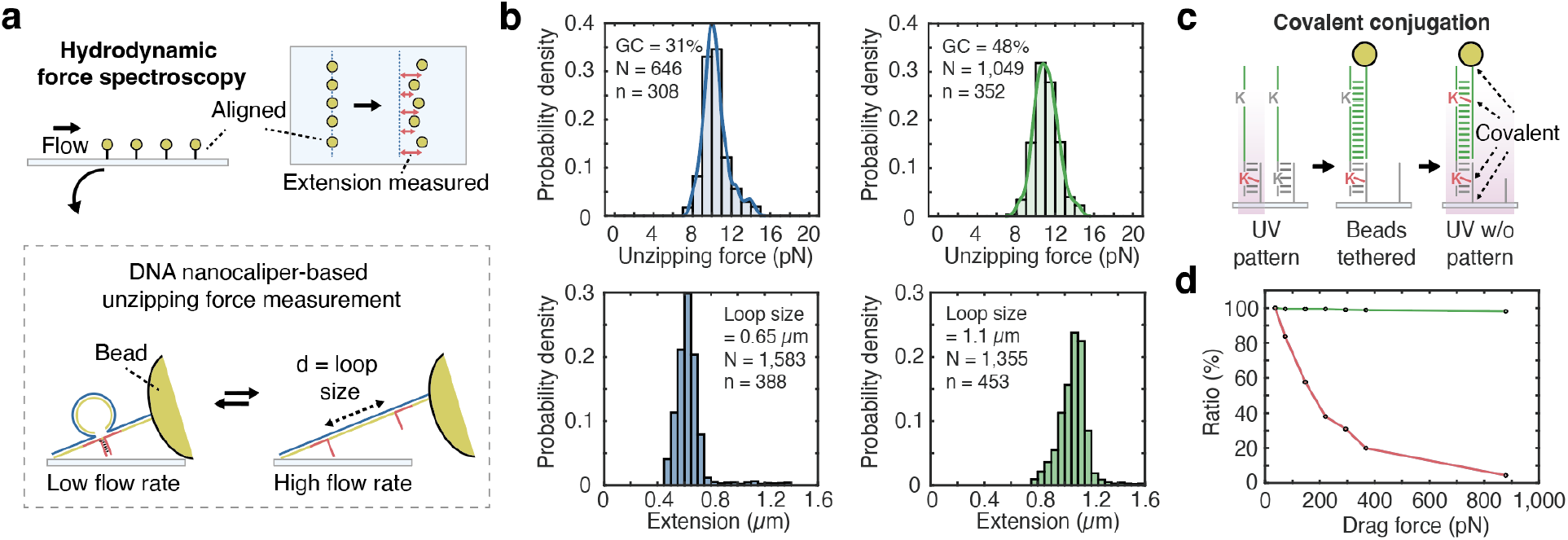
Hydrodynamic force spectroscopy conducted on patterned surfaces. (a) The positions of patterned beads are monitored during the application of hydrodynamic force. The same DNA nanoswitch constructs used in the magnetic tweezer experiments are employed. (b) Histograms of unzipping force with GC content of 31% (top left), and 48% (top right) measured using a 0.65 µm loop-sized construct are shown. Histograms of extension of DNA nanoswitch in X axis due to unzipping with loop sizes of 0.65 µm (bottom left) and 1.1 µm (bottom right) are shown. N represents the total number of measurements collected across n molecules. (c) Beads are patterned solely with covalent bonds by introducing two CNVK nucleosides in the patterning oligos. Following beads patterning by hybridization of functional region, UV is applied to the entire coverslip to covalently crosslink tethered beads. (d) The ratio of remaining beads on the patterned surface is measured by increasing the hydrodynamic force.

### Bead patterning with covalent bonds for high force application

Hydrodynamic force spectroscopy enables a relatively wide range of forces to be applied to single molecules, with a higher maximum force than typical magnetic tweezers systems. However, the maximum usable force is limited by the strength of the interactions tethering the beads to the surface—in particular, non-covalent interactions may simply not survive. To address this challenge, we adapted our pattering method to tether beads covalently by introducing two CNVK nucleosides to the patterning oligos (Fig. 5c). The bead patterning procedure is essentially the same as before, except for the introduction of a CNVK nucleoside in the functional sequence of the patterning oligo, and the usage of azide-coated magnetic beads. The beads with a diameter of 3.06 µm and DBCO-modified oligos were incubated to functionalize beads with oligos through copper-free click chemistry, and then functionalized beads were incubated onto the patterned flow cell to arrange them through oligo hybridization.

Following beads arrangement, UV was illuminated through a handheld UV lamp to the whole flow cell to covalently crosslink the oligos patterned on the flow cell with those functionalized on the beads via CNVK. Beads were organized in a square lattice pattern with 15.3 µm spacing, and the ratio of remaining beads under distinct hydrodynamic force application was investigated.

To evaluate the effectiveness of our covalent tethering approach, we performed hydrodynamic force spectroscopy experiments at high forces, comparing the covalently patterned surfaces to the non-covalently patterned surfaces. We applied the flow to both surfaces as a step function with 7 distinct flow rates, increasing sequentially from low to high, each maintained for 5 seconds. Drag force applied to beads is calculated by considering the bead size and shear rate (Supplementary Note 1). With the non-covalent approach, less than 40% of the tethered beads remained after application of a drag force of 220 pN per bead, corresponding to a flow rate of 300 µl/min. In contrast, the beads that were tethered covalently survived much higher forces, with 98% of the beads (n = 653) remaining even under application of a maximum drag force of 880 pN per bead, corresponding to a flow rate of 1,200 µl/min.

## Discussion

In this article, we proposed a light-guided molecular patterning method capable of precisely arranging biomolecules with multiple identities and beads in arbitrary patterns without the need for lithographic instruments. As a demonstration, oligos with unique sequences were precisely positioned and identified by tethering them to beads coated with distinct fluorophores. As an application, we showed how our method can enable multiplexed single-molecule force spectroscopy measurements, carrying out bond rupture experiments with both magnetic force and hydrodynamic force on patterned flow cells. By combining this assay with nanoengineered DNA nanoswitch constructs, we successfully measured the unzipping forces of oligos of different sequences. Uniquely, our approach can be used to covalently tether single molecules to the surface with high precision, enabling multiplexed experiments to be conducted at higher forces and throughput.

For comparison, previous approaches have demonstrated the ability to track multiple molecules by randomly positioning them on surfaces, either through physisorption with nitrocellulose^11^ or via covalent interactions, such as N-Hydroxysuccinimide (NHS) ester conjugation or click chemistry^38^. However, the lack of precise control over spatial positioning can limit throughput and the ability to perform spatially resolved experiments with multiple molecular identities. More recently, a soft lithography-based approach has been proposed to pattern the surfaces with proteins, enabling the precise arrangement of oligos using a polydimethylsiloxane (PDMS) stamp^18^. While effective in increasing throughput, this approach relies on non-covalent attachment via adsorption through a stamp and requires a master mold to be prepared through photolithography for each pattern, limiting its applicability to high applied forces and making it a challenge to patterning with multiple molecular identities. In contrast, our light-guided approach enables fabrication of arbitrary molecular patterns with diverse molecular identities by simply manipulating a DMD within a few seconds, and is well-suited for high force application. Moreover, crosslinked CNVK nucleoside can be split under irradiation of 312 nm UV^19^, allowing for potential resetting or reusing of specific regions of the flow cell multiple times. Our method utilizes a commercially available DMD (Polygon 400, Mightex Inc.) that can be easily attached to microscope, and it can be used to pattern biomolecules through conjugation to oligos for various purposes such as proteins for cell patterning and nanoparticles for photonics applications.

Our light-guided patterning method can enhance the throughput of biomolecular assays by enabling the rapid and precise patterning of different molecular species, thereby increasing the number of trackable molecules— improvements that should have a particularly large impact on single-molecule approaches. By virtue of its programmability, versatility—facilitating both covalent and non-covalent attachments—and its speed and accessibility, we expect this technology to impact diverse research fields including biophysics, nanotechnology, molecular diagnostics, and biophotonics.

## Acknowledgments

This work was supported by the National Institute of General Medical Sciences (NIGMS; R35GM119537) (W.P.W.) and the International Human Frontier Science Program Organization (LT0005/2023-C) (H.C.). We thank Peng Yin and Ninning Liu at the Wyss Institute at Harvard University for their assistance with DMD instrument, and J. Zsolt Terdik, Prakash Shrestha, and Darren Yang at Boston Children’s Hospital for helpful discussions.

## Author Contributions

H.C., and W.P.W. conceptualized and designed the experiments. H.C. and A.W. conducted single-molecule force spectroscopy experiments. H.C., A.W., and W.P.W. participated in data analysis and discussion. H.C., and W.P.W wrote the original draft and edited the manuscript.

## Methods

### Oligo design and DNA nanoswitch construct preparation

The 3’ end of the 17 nt base oligo is modified with DBCO-PEG13 (Polyethylene glycol linker) to attach it to the azide coverslip (104-00-626, PolyAn) and passivate the surface. Each patterning oligo consists of a 55 nt construct hybridization region as well as a 14 nt base oligo hybridization region that includes a CNVK nucleoside to crosslink it upon UV illumination (Supplementary Fig. 2 and Supplementary Table 1). The DNA nanoswitch construct for force spectroscopy is synthesized with circular M13mp18 ssDNA (N4040S, New England Biolabs). To linearize the circular DNA, 10 µl of 110 nM M13mp18 ssDNA, 7 µl of 10 µM 20 nt of cutting strand (Supplementary Table 1.) that hybridizes to the cutting region in circular DNA, and 2 µl of 10X rCutSmart buffer were annealed for 20 minutes by heating to 90°C and slowly cooling down to 50°C at a rate of 30 seconds /°C. Next, 1 µl of BtsCI restriction enzyme (R0647S, New England Biolabs) is added followed by incubation at 50°C for an hour and inactivation at 80°C for 20 minutes. To fabricate the dsDNA linear construct, linearized M13mp19 ssDNA at 1nM concentration was incubated with 5 nM of backbone oligos 1–119 (Supplementary Data 1), and 2 nM of backbone-biotin oligo by heating to 90°C and slowly cooling down to 80°C at a rate of 1 minute/°C and 80°C to 20°C at a rate of 6 min/°C. To fabricate the construct with a loop size of 1.1 µm, backbone oligos 116 and 117 were replaced with unzipping oligos with overhangs (Supplementary Table 1). To fabricate the construct with a 0.65 µm loop, backbone oligos 118 and 119 were replaced. Fabricated constructs were analyzed with 0.7% agarose gel, which was run at 100V for around 2 hours in a 4C room. Base oligo with DBCO-PEG13 modification and patterning oligos with CNVK modification were purchased from Gene Link and other oligos were purchased from IDT (Integrated DNA Technologies) unless otherwise specified.

### Flow cell preparation and oligo patterning

Buffer A (PBS, pH 7.4), buffer B (PBS, 1% Tween 20, pH 7.4), buffer C (PBS, 1% Tween 20, 500 mM NaCl, 0.01% (w/v) blocking reagent; 11096176001, Roche), buffer D (PBS, 1% Tween 20, 0.01% (w/v) blocking reagent), and blocking buffer (PBS, 1% Tween 20, 1% (w/v) blocking reagent) were prepared for the experiments. Flow cells were fabricated by sandwiching glass slides (48300-037, VWR International) and azide-coated coverslips with double-sided tape^39^. A flow chamber with channel dimensions of 1.96 mm width, 26.46 mm length, and 100 µm height was created by cutting double-sided tape using a cutter plotter (CE7000-60, Graphtec), prior to assembly.

Glass slides were drilled using a diamond drill bit with a diameter of 0.75 mm (415601, SFJS) to make inlets and outlets. The channel was incubated with 100 µM of DBCO-PEG13 modified base oligo in buffer A for 18 hours. After washing the channel three times with buffer B, the channel was incubated with the blocking buffer for 30 minutes twice followed by washing with buffer C. 100 nM of patterning oligo in buffer C was incubated for 10 minutes and the desired pattern of 365 nm UV was illuminated following the previously reported method^21^. Briefly, 365 nm UV (BLS-Series 50W emitter, Mightex) was illuminated through a 20X objective with numerical aperture (NA) of 0.5 (CFI Plan Fluor, Nikon) at a power setting of 4% for 2 seconds. UV power can be adjusted based on the number of on-target beads, and the number of singly tethered beads as increased power will increase the crosslinking efficiency^30^. Patterning oligos that are not crosslinked to the base oligos are washed away by incubating with 40% formamide for 3 minutes followed by washing with buffer D. Flow cells were stored with buffer D at 4°C or within a vacuum chamber and were incubated with blocking buffer for an hour before each experiment.

### Bead conjugation to each patterned spot

M270 Dynabeads (65305, ThermoFisher Scientific) were used for magnetic tweezers and hydrodynamic force spectroscopy experiments. To prepare the beads for experiments of non-covalently attached beads, 10 µl of 10mg/ml beads were washed with 1X Binding and Wash (B&W) buffer (50 mM sodium phosphate, 300 mM NaCl, and 1% Tween20) for three times followed by incubation with 100 µl of 10 nM biotinylated functional oligo in 1X B&W buffer (Supplementary Table 1). Next, the beads were washed with buffer B three times, and the surface of the beads was blocked by incubation with 50 µl of blocking buffer for 1 hour. Beads were washed with buffer B before usage. The flow cell and 1 ml gastight syringe (1001-TLL, Hamilton Company) were connected through peek tubing (1569, P628, F-247, and XP-235, IDEX Corporation), and beads were added to the flow cell at a flow rate of 20 µl/min using a syringe pump (70-4504, Harvard Apparatus). To functionalize the surface of the flow cell, 1 nM of DNA nanoswitch construct was incubated with the flow cell for 3 hours at room temperature followed by wash and bead incubation steps. For the covalent conjugation of beads, azide magnetic beads (CBPMC01c, Bangs Laboratory Inc.) and 55 nt DBCO-modified functional oligo were used. Washed azide beads (100 µg) were incubated with 100 µl of 100 nM DBCO oligos for two hours. Beads were washed with buffer B three times and incubated with blocking buffer for an hour before usage. After bead assembly through hybridization of the functional sequence, the flow cell was carefully moved to the top of a handheld 6W UV lamp (97620-22, Cole-Parmer) and illuminated with 365 nm UV light for 30 minutes.

