## Supplementary Materials for "Light-guided molecular patterning for programmable multiplexed single-molecule manipulation"

**This file includes:**

Supplementary Note 1

Supplementary Figures 1 – 15

Supplementary Tables 1

**Supplementary Note 1.** Magnetic tweezers and flow cell force calibration & experiments

Magnetic force applied to beads was calculated using the equipartition theorem[1–3] where $F = \frac{k_B T L}{<X^2>}$, $k_B$ is the Boltzmann constant, $T$ is absolute temperature, $L$ is extension of the tether, and $<X^2>$ is the mean square displacement from the mean position along the x-axis, which was parallel to magnetic field. To determine the force calibration curve for different magnet heights, bar magnets with a 1 mm gap were placed at a variety of different Z positions (0.5 mm, 1 mm, 2 mm, and 5 mm) from the glass slide while $L$ and $<X^2>$ were measured[4]. The distance between the coverslip surface and the top side of the glass slide was 1.3 mm since the channel height was 0.1 mm and the glass thickness was 1.2 mm. To interpolate between different magnet positions, the force vs magnet position data was fitted to an exponential curve. To measure DNA unzipping events, the magnet was moved from a position of 10 mm to a position of 0.3 mm with a magnet speed of 1 mm/s; images were taken with a frame rate of 50 frame per second (fps). For the magnetic tweezer experiments, a 60X objective with an NA of 1.27 (Plan Apo IR, water immersion, Nikon) was used.

For the calibration of the hydrodynamic force under flow, we calculated the geometry of the system and the tension on the DNA construct at different flow rates using a previously reported method[5] based on fluid dynamics theory. Using force balance and torque balance and the hydrodynamics of laminar flow near a surface, one can derive Equations 1 and 2, where a = 1.7, the ratio of torque to drag force R = 0.37r, with r as the bead radius, $\eta$ is the flow viscosity, which we set to $\eta = 10^{-3} Ns/m^2$, $F_T$ is the tension, $F_{drag}$ is the drag force applied to the bead, v is the flow velocity, $\alpha$ is the tether angle relative to the coverslip, and $\kappa$ is the angle between the imaginary line connecting the point where the bead touches the surface and the DNA-bead attachment point, and the vertical line passing through the bead's center (Supplementary Fig. 1). The flow velocity v was calculated as $r\gamma$, where the shear rate $\gamma = \frac{6Q}{wh^2}$, Here Q is the volumetric flow rate, w = 1.9 mm is the channel width, and h = 100 µm is the channel height, which simplifies the equation to v (µm/s) = 7.15Q (µl/min) when 2.8 µm beads are used. Equation 1, and 2 are derived from force and torque balance, while equation 3 relating $x, \alpha, and\ \kappa$ is derived from the geometry of the system (Supplementary Fig. 1). Equation 4, which relates $x$ and $\alpha$ is obtained by combining Equations 1–3. Additionally, the bead center to center distance ($D_{cc}$) during flow infusion and withdrawal is experimentally measured, and Equation 5 is derived based on the system's geometry. To measure $D_{cc}$ for the DNA nanoswitch construct in the looped conformation, we supplemented the buffer with 20 mM of $MgCl_2$ to increase the unzipping force and used higher loading rates. The flow infusion rate was ramped from 0 µl/min to 50 µl/min for 5 seconds, followed by a flow withdrawal at a constant 50 µl/min flow rate. The $D_{cc}$ was calculated under the assumption that the extension profile of a single construct should be symmetric between flow infusion and withdrawal.

$$Force\ balance: F_T cos(\alpha) = F_{drag} = a6\pi\eta rv \quad (1)$$

$$Torque\ balance: F_T cos(2\kappa - \alpha) = F_{drag} R \quad (2)$$

$$From\ geometry: xsin(\alpha) = r\big(1 - cos(\pi - 2\kappa)\big) \quad (3)$$

$$\tan(\alpha) = \frac{1 + R - \frac{xsin(\alpha)}{r}}{\sqrt{(\frac{xsin(\alpha)}{r})^2 + \frac{2xsin(\alpha)}{r}}} \quad (4)$$

$$From\ geometry: D_{cc} = 2(x(cos(\alpha) + rsin(2\kappa)) \quad (5)$$

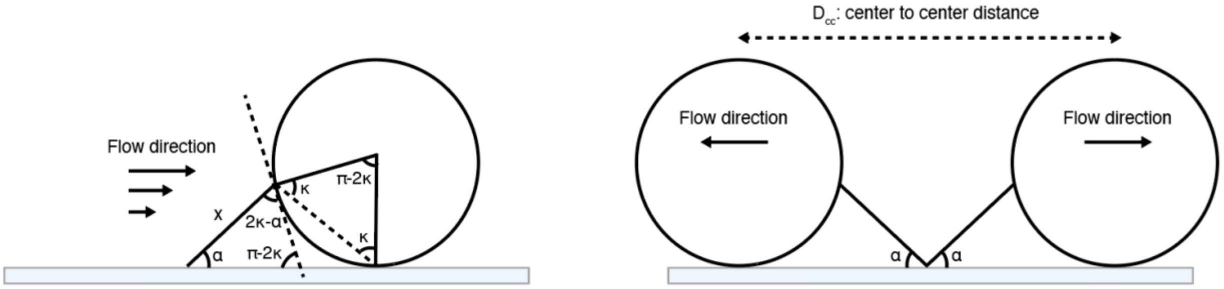


**Supplementary Figure 1.** Geometry of a bead tethered to a DNA construct under flow conditions[5].

As most of the unzipping was observed between the flow rate of 25 µl/min and 30 µl/min, we calculated the angles and $x$ by numerically solving the equations at these two flow rates using MATLAB. For the constructs with a loop size of 0.65 µm, we found $\alpha = 32.71°, \kappa = 57.34°,\ x = 1.51\ \mu m, and\ F_T = 9.53\ pN$ when the flow rate was 25 µl/min and $\alpha = 32.59°, \kappa = 57.28°,\ x = 1.52\ \mu m, and\ F_T = 11.4\ pN$ when the flow rate was 30 µl/min. For the constructs with a loop size of 1.1 µm, we found $\alpha = 36.45°, \kappa = 59.27°,\ x = 1.23\ \mu m,\ and\ F_T = 9.97\ pN$ when the flow rate was 25 µl/min and $\alpha = 36.45°, \kappa = 59.27°, x = 1.23\ \mu m, and\ F_T = 12\ pN$ when the flow rate was 30 µl/min. Next, to interpolate the force as a function of flow rate, we assumed that the force is linearly correlated to the flow velocity within this range of flow rates as the calculated change in angle was small between the flow rates of 25 µl/min and 30 µl/min, and since $F_T = \frac{F_{drag}}{\cos(\alpha)}$ where drag force is proportional to flow velocity.

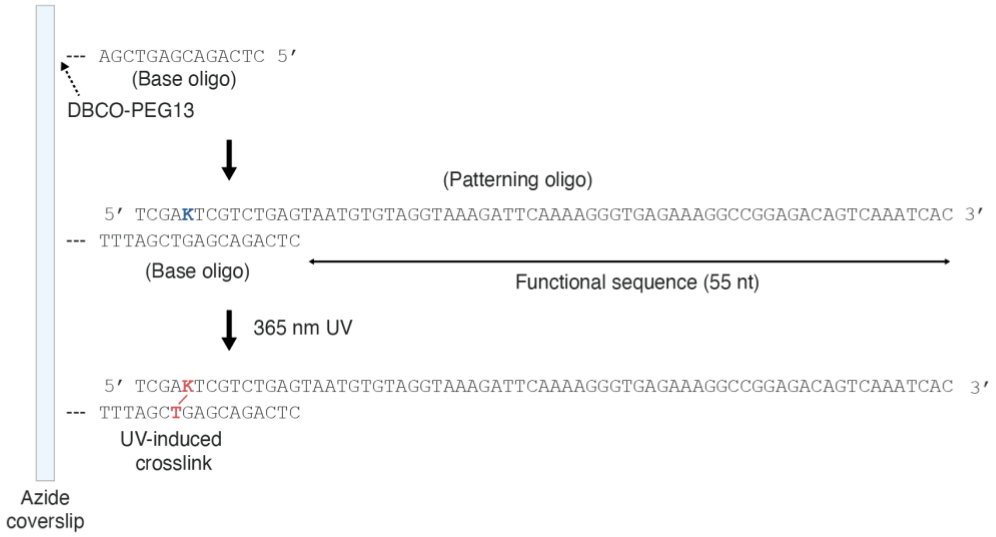


**Supplementary Figure 2.** Base oligo and patterning oligo sequence design.

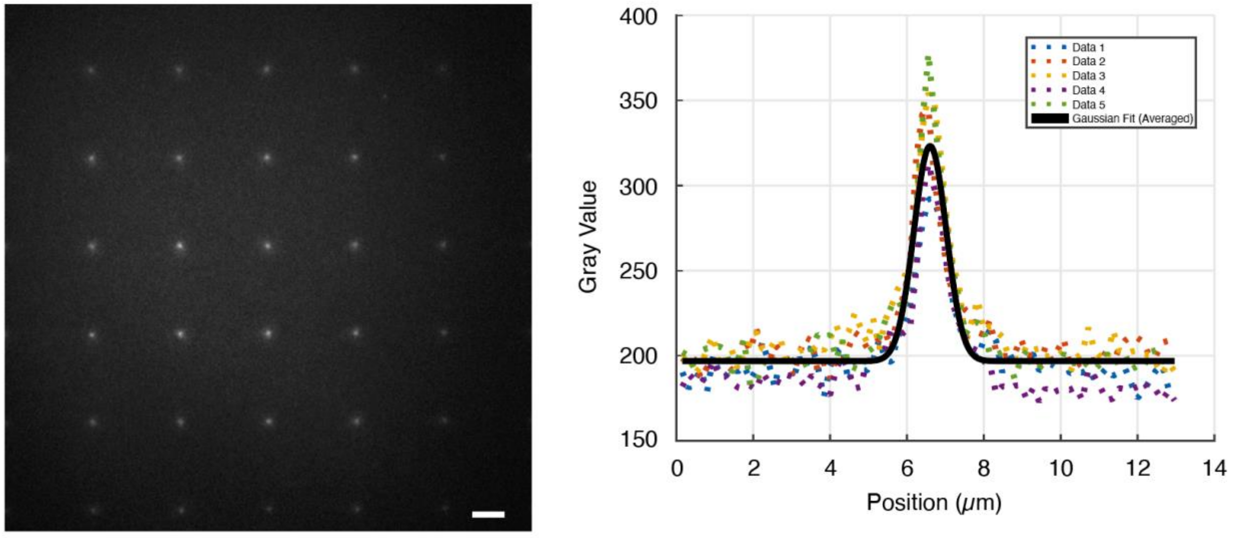


**Supplementary Figure 3.** A square array with each spot patterned using a single micromirror was analyzed by hybridizing fluorescent oligos. The fluorescence intensity from five spots was measured and fitted to a Gaussian curve, resulting in a full width at half maximum of 992 nm. Scale bar, 5 μm.

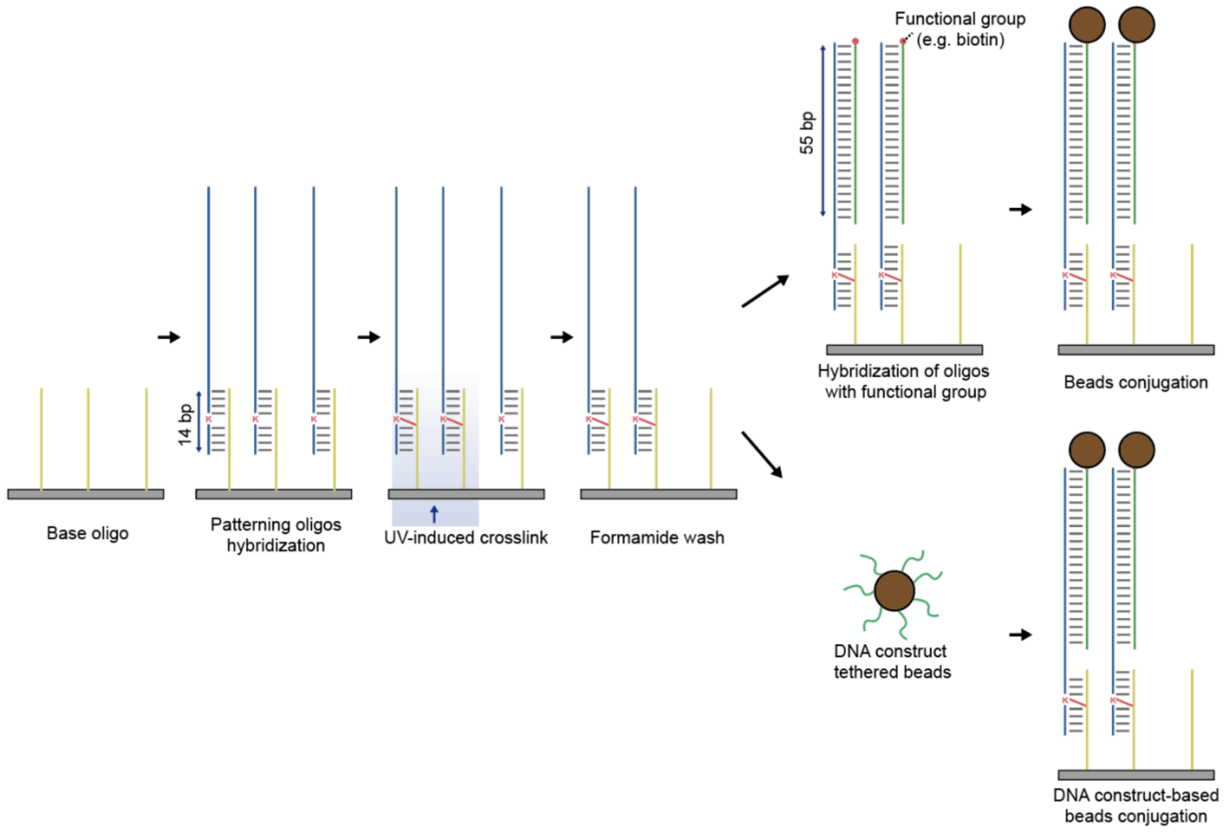


**Supplementary Figure 4.** The DNA construct connects the patterning oligo and beads. It can hybridize with patterning oligos to tether beads or be attached to the beads first, followed by hybridization with the patterning oligos.

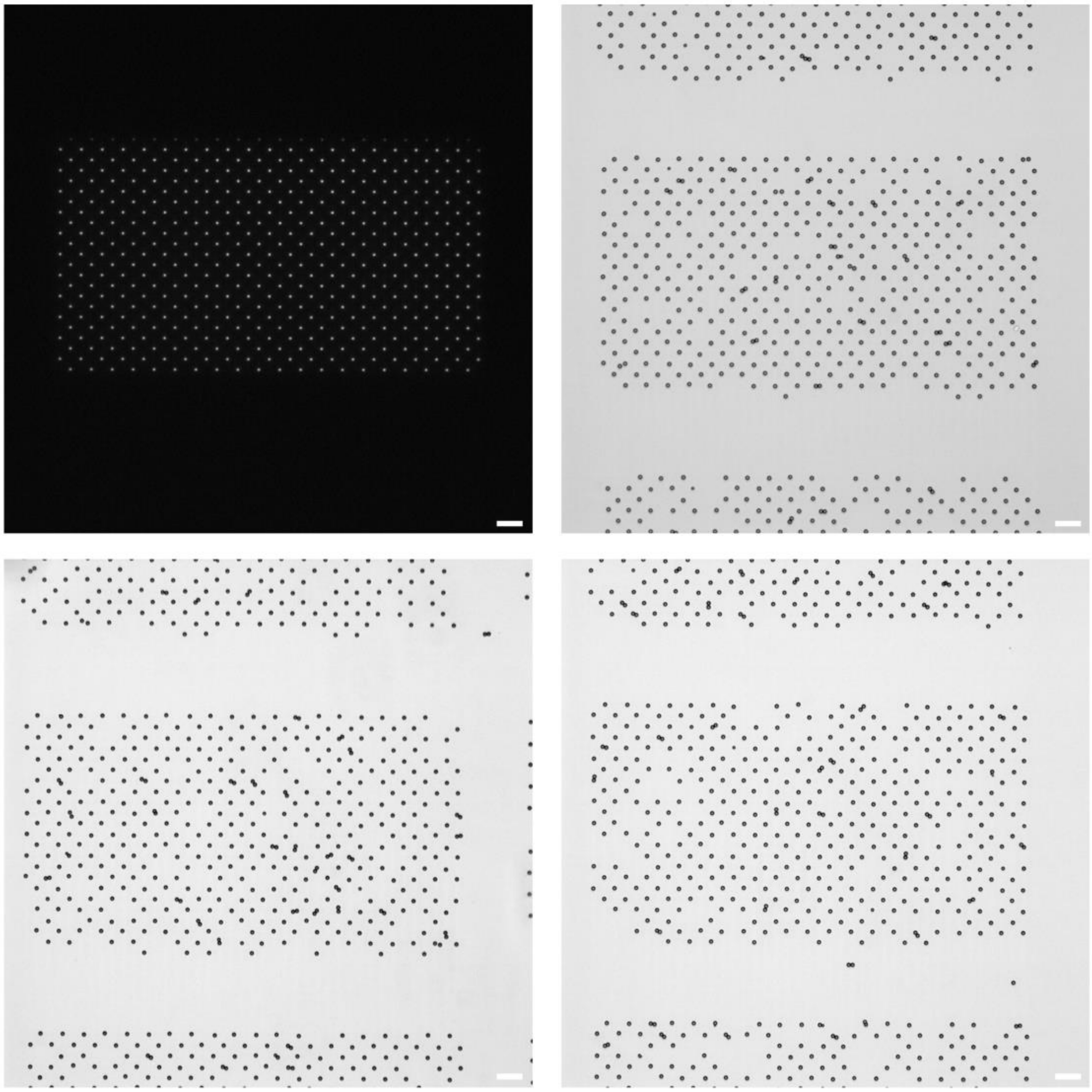

**Supplementary Figure 5.** Beads are arranged in square array lattice with intended distance of 11.5 µm. Left top image shows UV illumination pattern on blank coverslip. Scale bar, 20 µm.

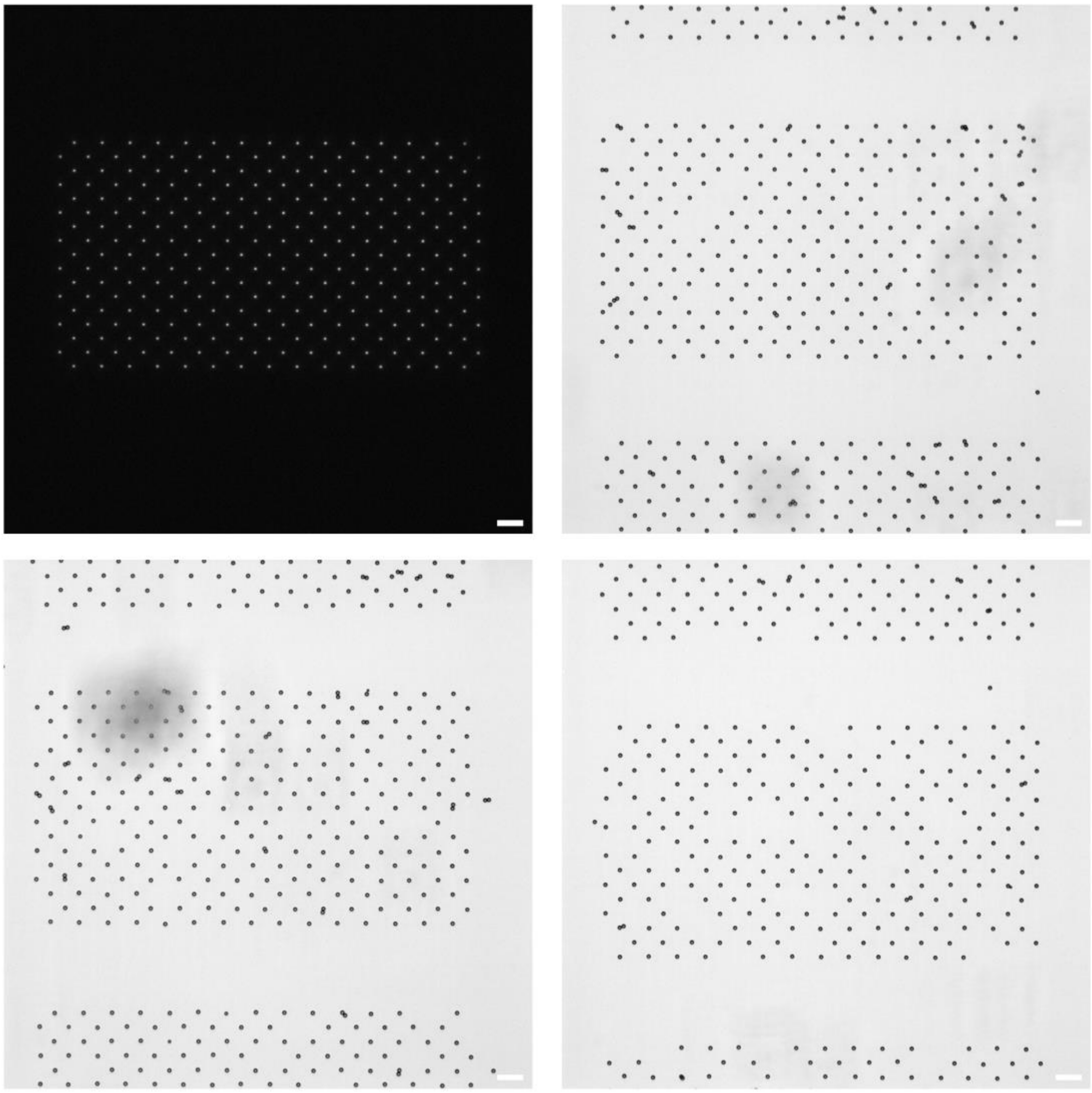

**Supplementary Figure 6.** Beads are arranged in square array lattice with intended distance of 15.3 µm. Left top image shows UV illumination pattern on blank coverslip. Scale bar, 20 µm.

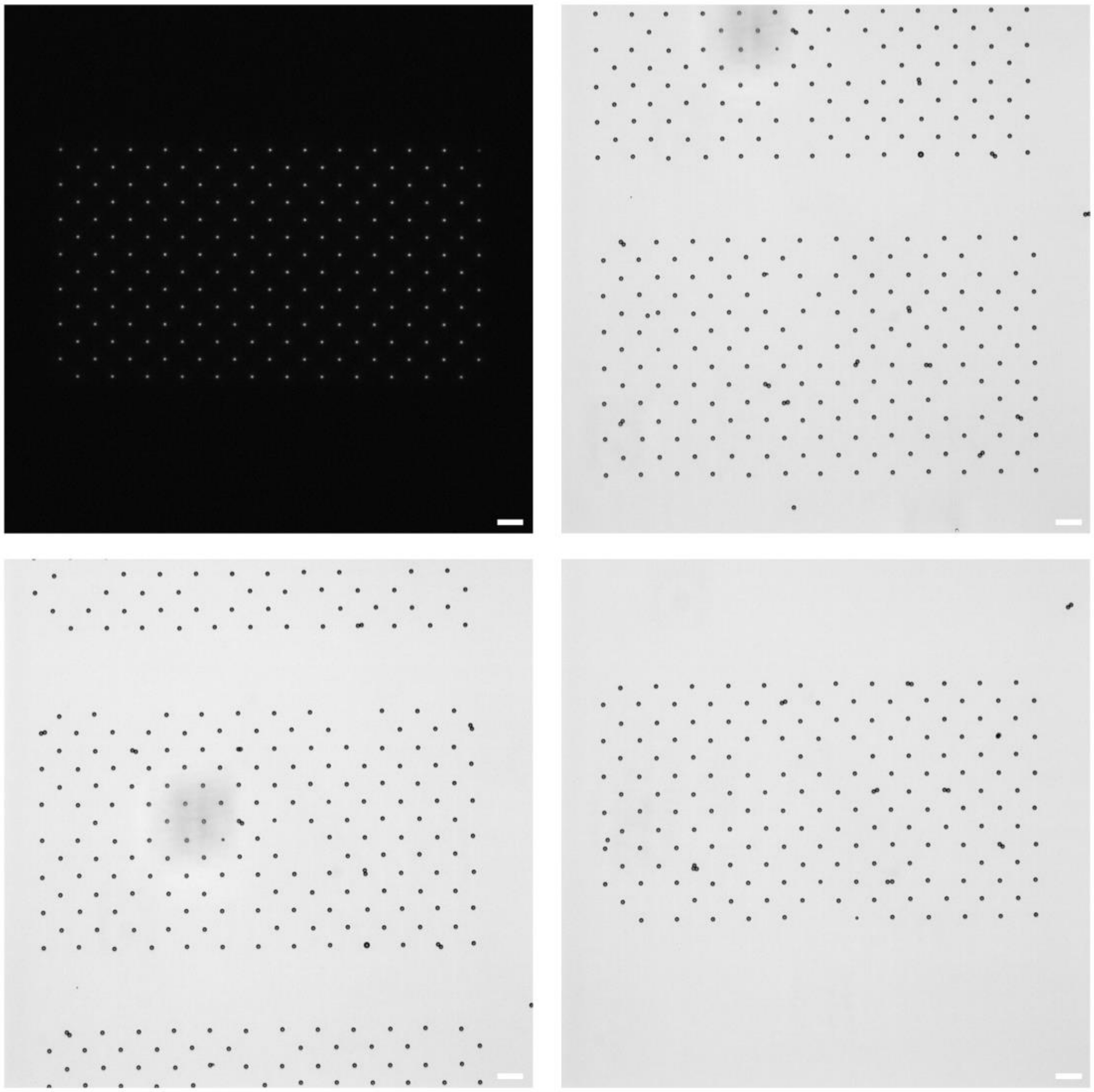

**Supplementary Figure 7.** Beads are arranged in square array lattice with intended distance of 19.2 μm. Left top image shows UV illumination pattern on blank coverslip. Scale bar, 20 μm.

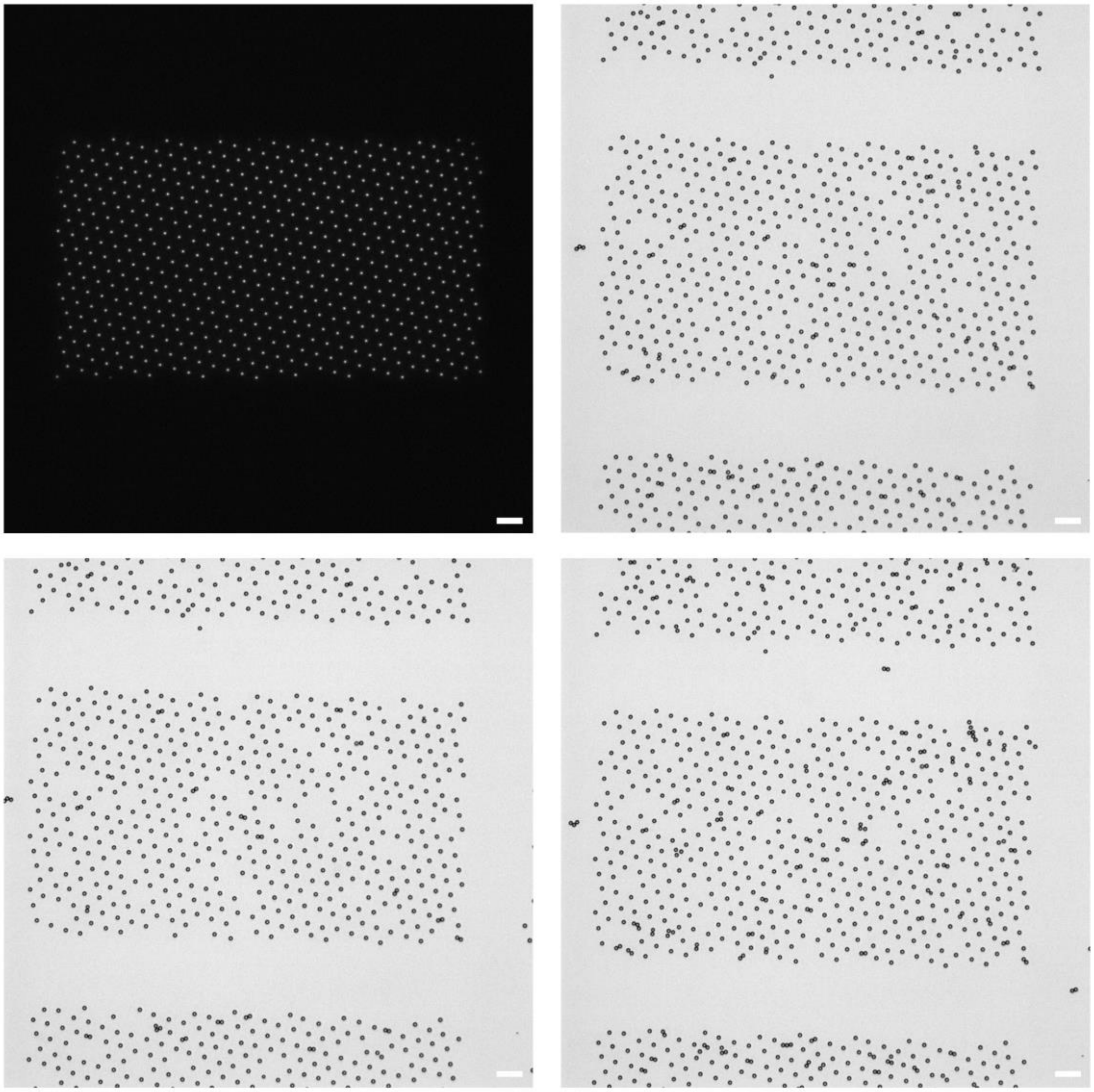

**Supplementary Figure 8.** Beads are arranged in hexagonal array lattice with intended distance of 11.5 µm. Left top image shows UV illumination pattern on blank coverslip. Scale bar, 20 µm.

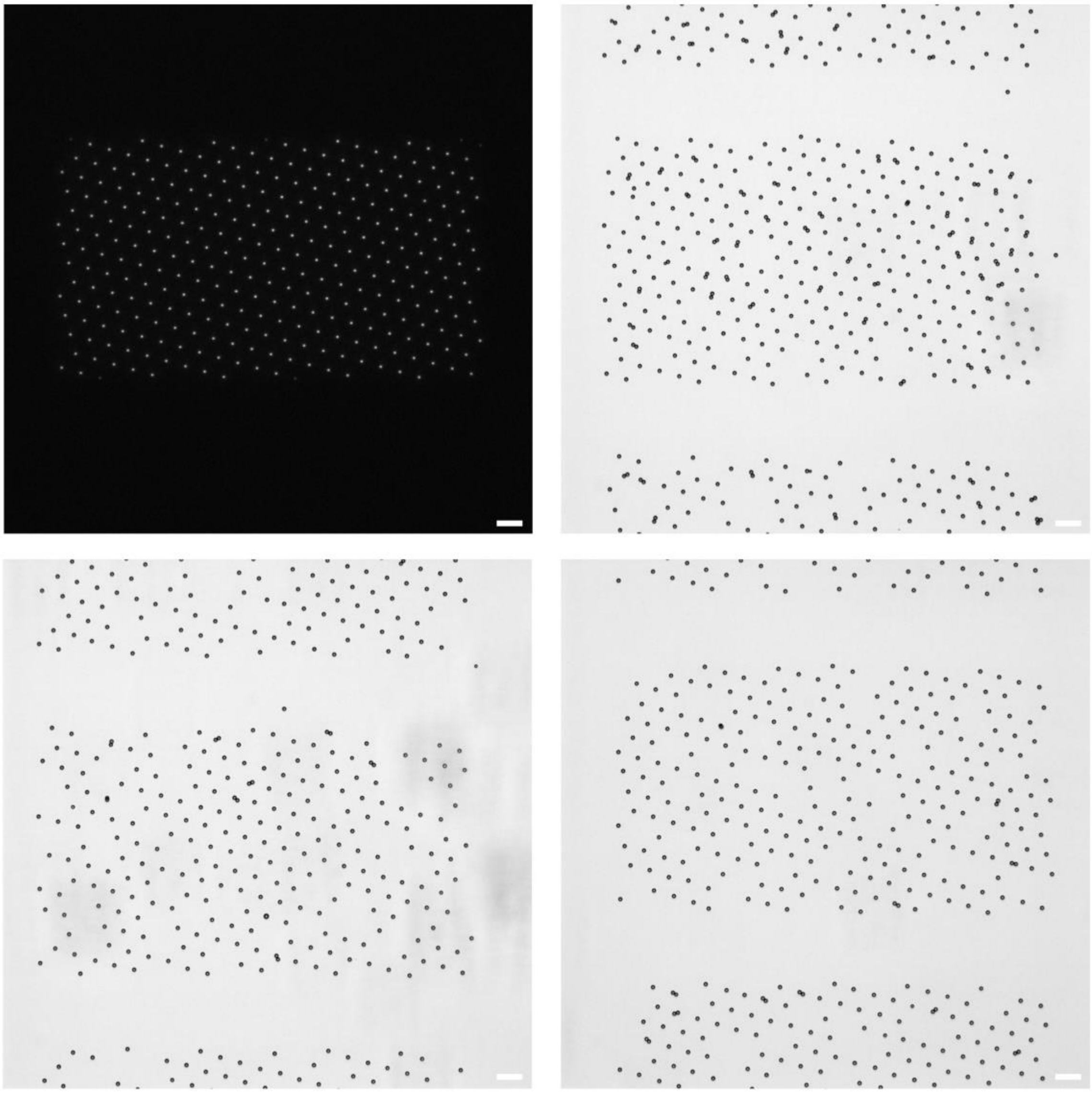

**Supplementary Figure 9.** Beads are arranged in hexagonal array lattice with intended distance of 15.3 μm. Left top image shows UV illumination pattern on blank coverslip. Scale bar, 20 μm.

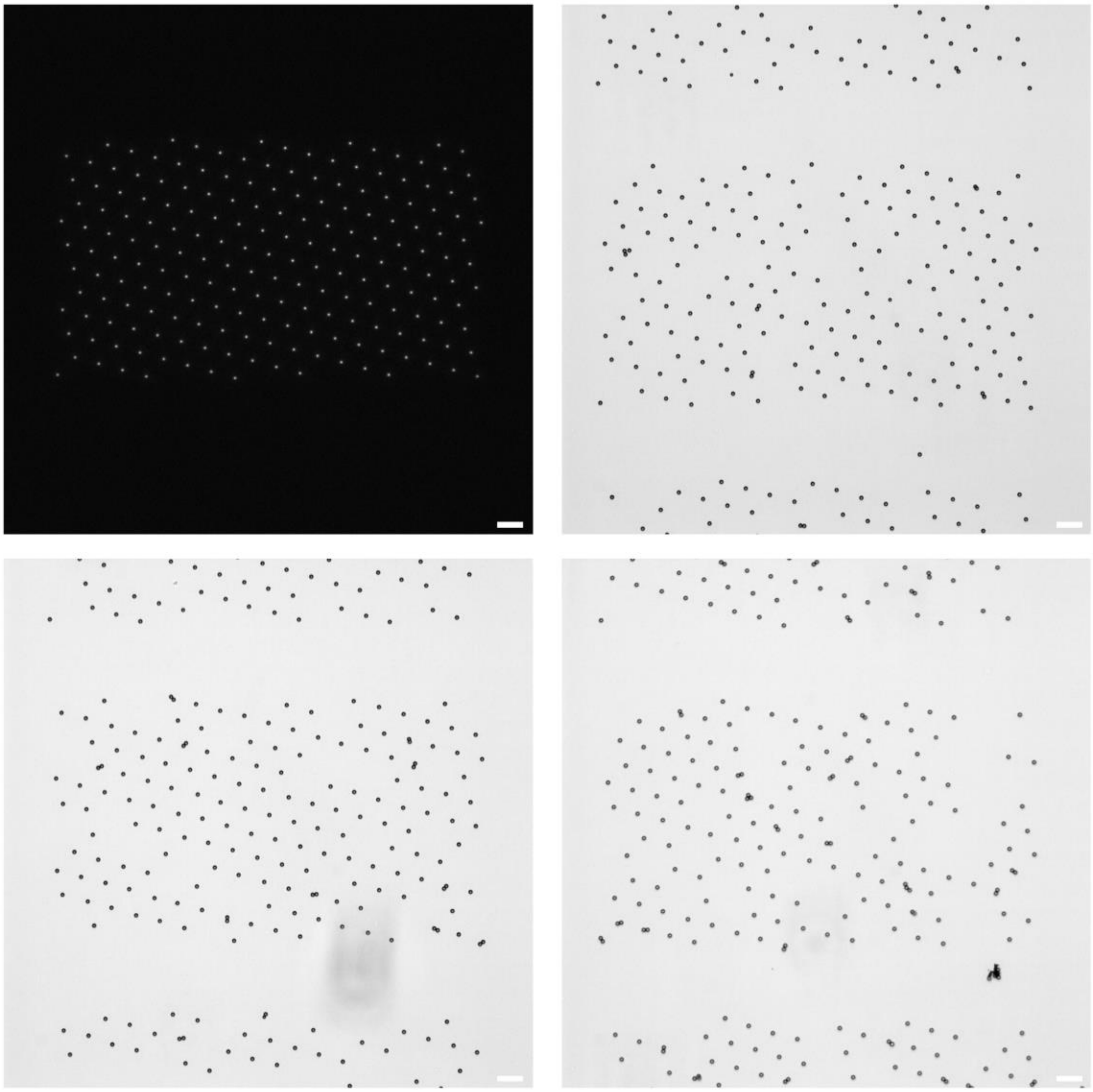

**Supplementary Figure 10.** Beads are arranged in hexagonal array lattice with intended distance of 19.2 µm. Left top image shows UV illumination pattern on blank coverslip. Scale bar, 20 µm.

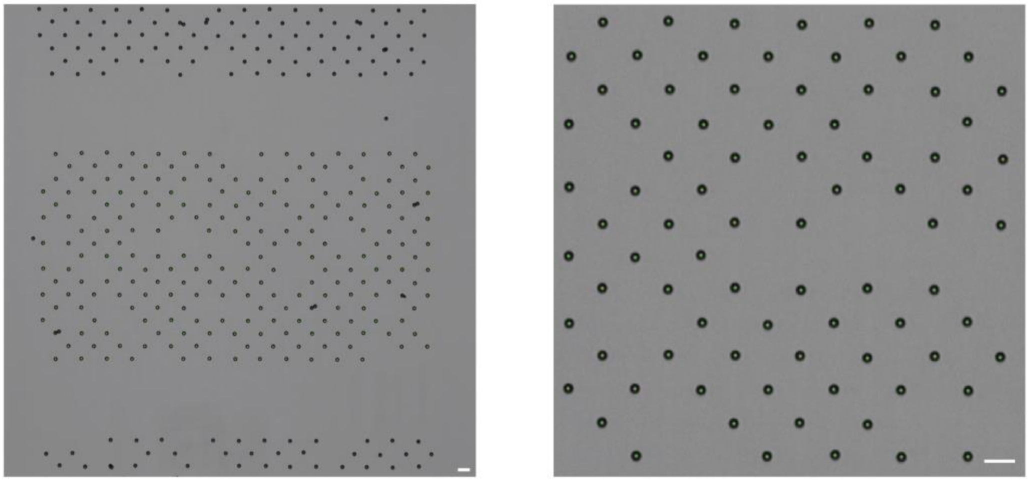

**Supplementary Figure 11.** Green asterisks show detected center of beads that were detected with edge detection algorithm using Sobel approximation. Example images of a bead array with a square lattice and 15.3 μm spacing are shown. Nonspecific and stuck beads are excluded by setting a region of interest and limiting the size of the detected circle. Scale bar, 10 μm.

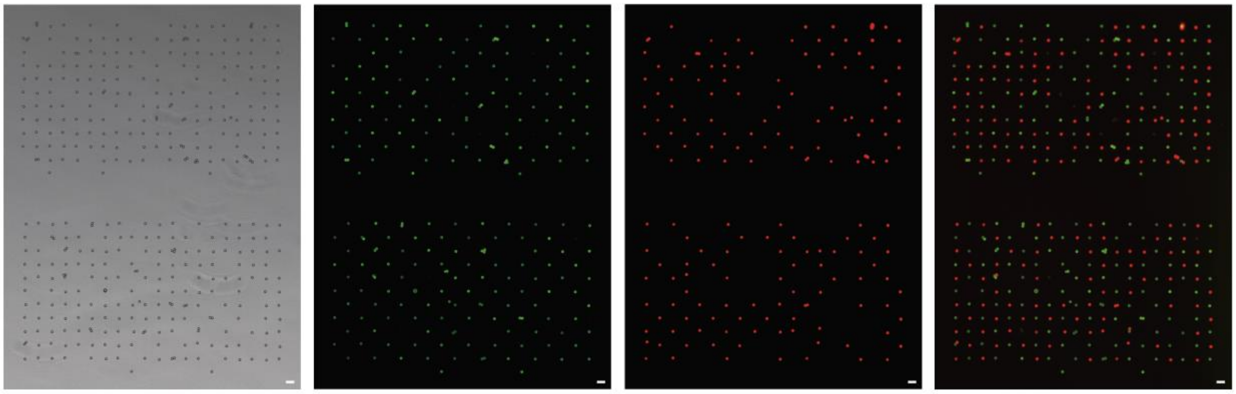

**Supplementary Figure 12.** A square bead array is fabricated with two distinct patterning oligos with different functional sequences. From left to right, the images taken with bright field, fluorescence channel with a Texas Red filter, fluorescence channel with a Cy5 filter, and the merged image are shown. Scale bar, 10 μm

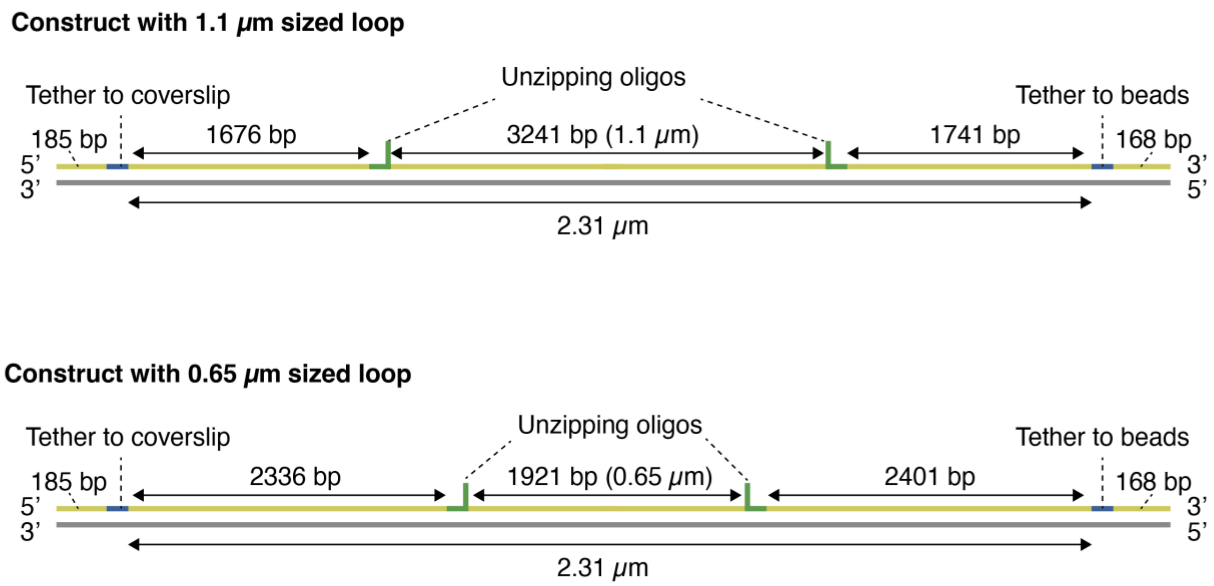


**Supplementary Figure 13.** DNA nanoswitch construct design for single-molecule force measurements. Oligo sequences are available in Supplementary Data 1 and Supplementary Table 1.

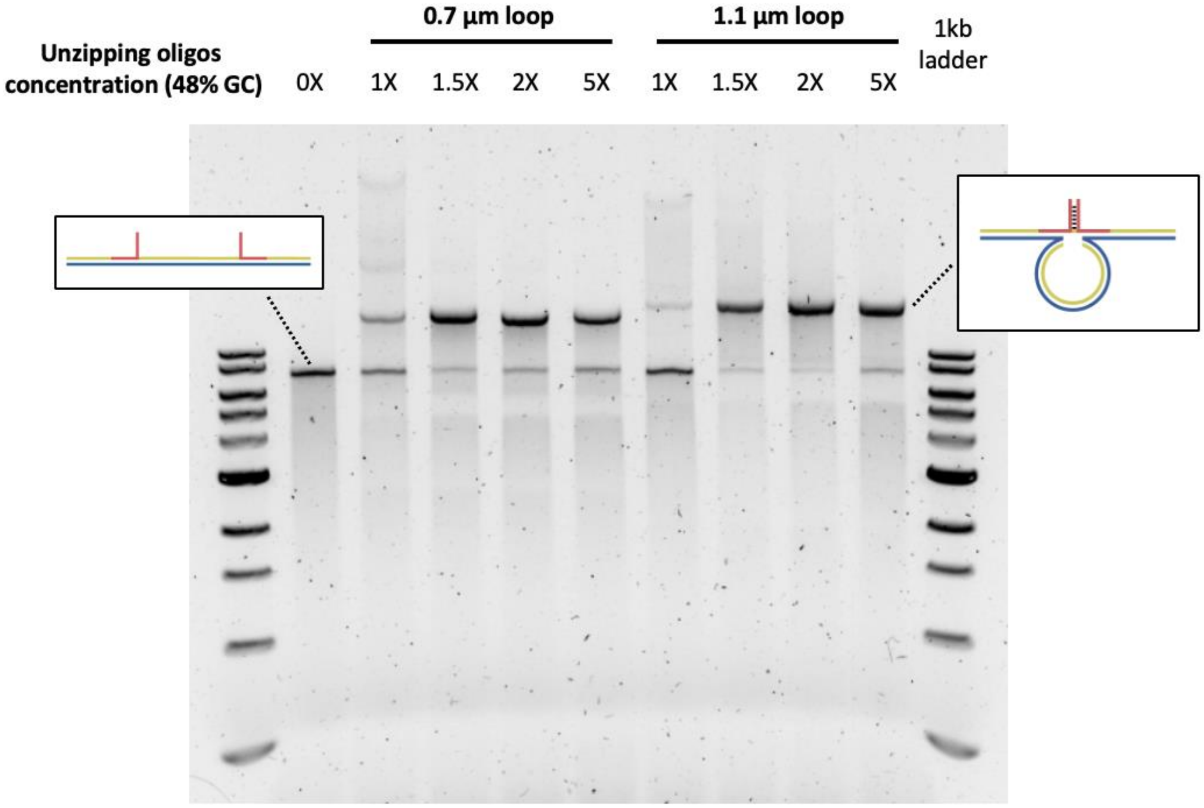


**Supplementary Figure 14.** 0.7% Agarose gel electrophoresis result of synthesized constructs. Final concentration of scaffold strand was 1 nM, and that of backbone oligos were 5 nM. Unzipping oligos with GC content of 48% were added at a concentration of 1 nM, 1.5 nM, 2 nM, and 5 nM and resulting looping ratio were 50.7%, 92.5%, 88.9%, and 80.2% respectively when the size of the loop was 0.7 µm. When the construct with 1.1 µm long loop were fabricated with unzipping oligos concentration of 1 nM, 1.5 nM, 2 nM, and 5 nM, the ratio of looped constructs were 13.2%, 95.2%, 97.5%, and 89.3%.

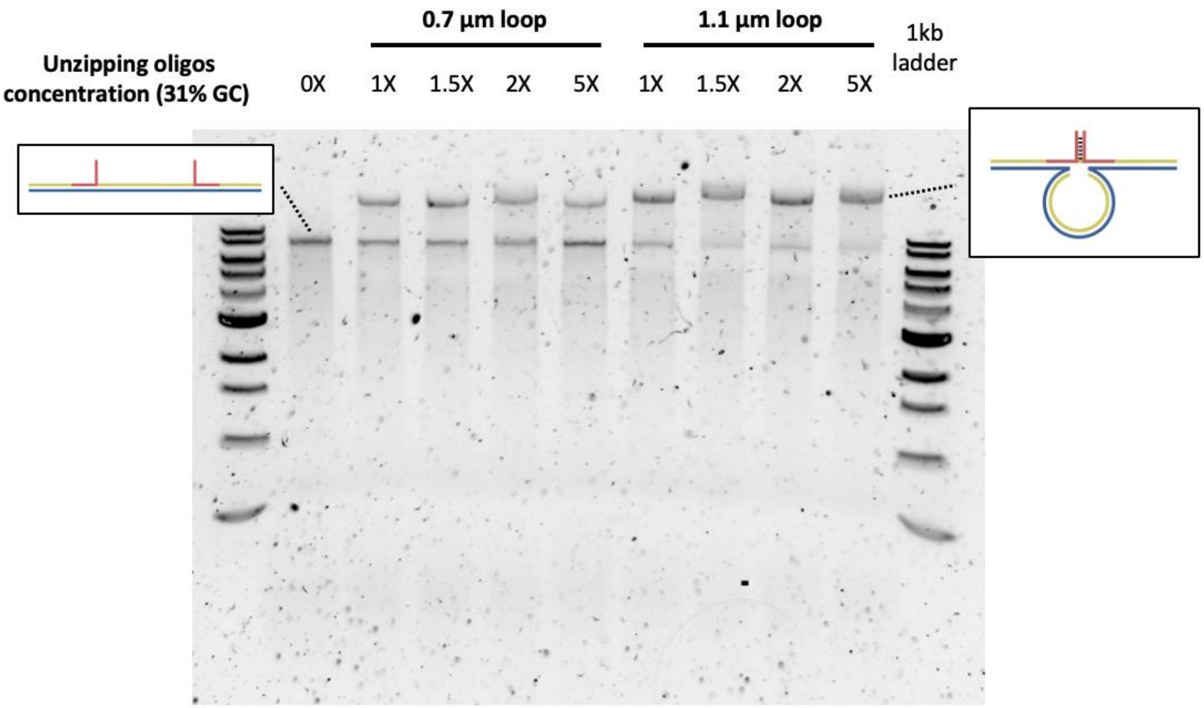


**Supplementary Figure 15.** 0.7% Agarose gel electrophoresis result of synthesized constructs. Final concentration of scaffold strand was 1 nM, and that of backbone oligos were 5 nM. Unzipping oligos with GC content of 31% were added at a concentration of 1 nM, 1.5 nM, 2 nM, and 5 nM and resulting looping ratio were 62%, 59.8%, 51.2%, and 40.9% respectively when the size of the loop was 0.7 µm. When the construct with 1.1 µm long loop were fabricated with unzipping oligos concentration of 1 nM, 1.5 nM, 2 nM, and 5 nM, the ratio of looped constructs were 75.8%, 83.3%, 84.6%, and 88.2%.

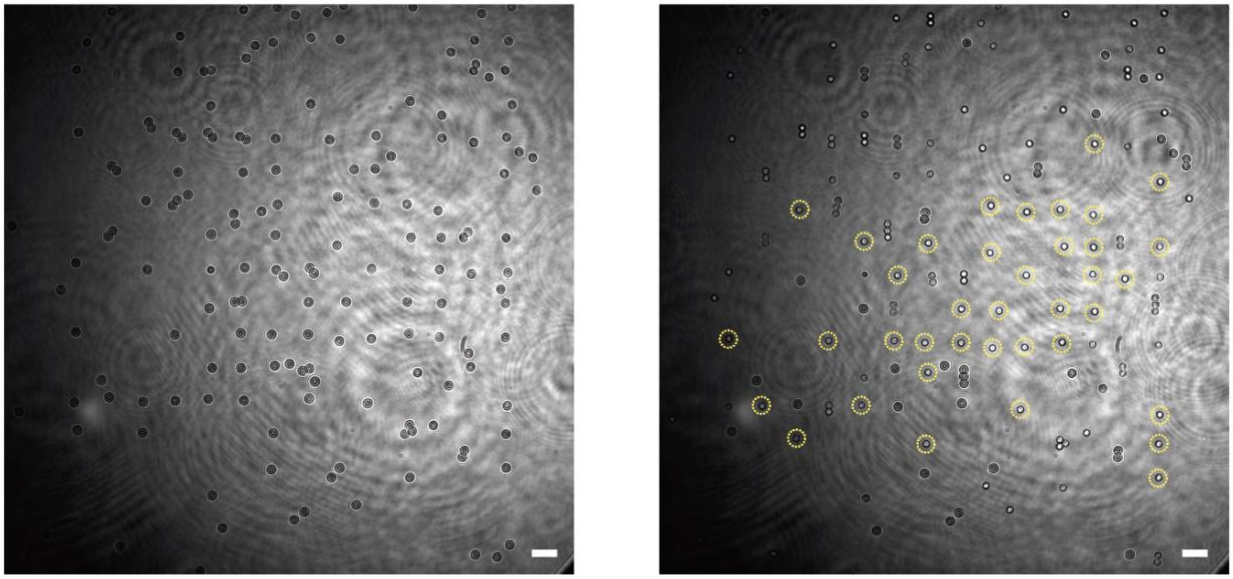

**Supplementary Figure 16.** Large field of view image of beads tethered with a 2.3 µm long DNA construct during a magnetic tweezer experiment. The left image was taken without magnetic force, and the right image was taken with a magnetic force of 15 pN. Circled beads indicate those tethered with a single construct.

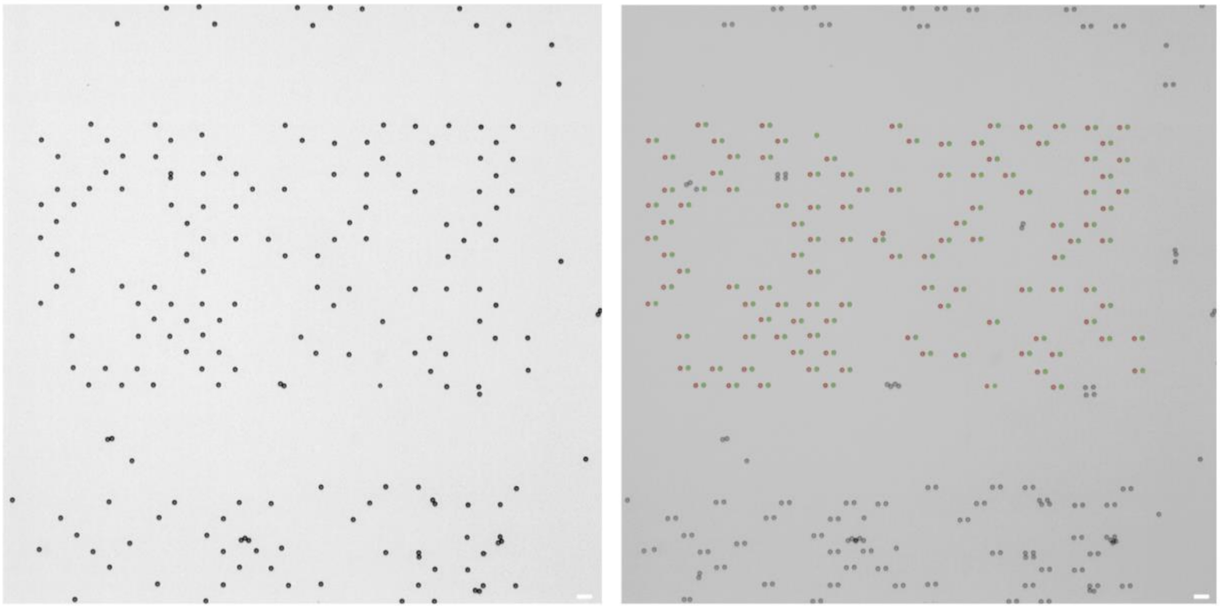

**Supplementary Figure 17.** Large field of view image of beads tethered with 2.3 µm long DNA construct. Right image shows merged image from flow infuse and withdrawal, showing the tether lengths. Beads were detected as red circle from the flow infuse, and green asterisk shows detected circle during flow withdrawal.

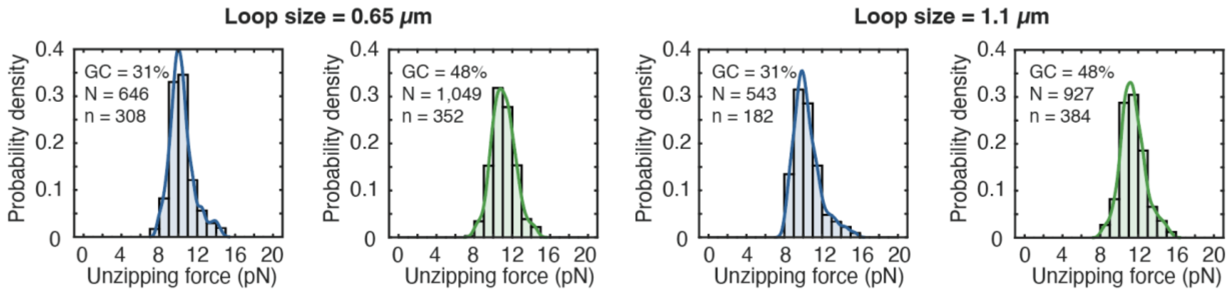


**Supplementary Figure 18.** Unzipping forces measured in a hydrodynamic experimental system, determined by loop sizes and the sequences of the unzipping oligonucleotides.

| | |
|---|---|
| Base oligo | CTCAGACGAGTCGATTT-[DBCO–PEG13]-3’ |
| Patterning oligo | TCGA[CNVK]TCGTCTGAGTAATGTGTAGGTAAAGATTCAAAAGGGTGAGAAAG<br>GCCGGAGACAGTCAAATCAC |
| Functional oligo | 5’ [Dual-biotin or amine]<br>TTTTTGTGATTTGACTGTCTCCGGCCTTTCTCACCCTTTTGAATCTTTACCTACA |
| Fluorescent oligo | 5’ [ATTO488]TTTTTGTGATTTGACTGTCTCCGGC |
| Cutting strand | CTACTAATAGTAGTAGCATTAACATCCAATAAATCATACA |
| 48% GC, 1.1 µm loop, unzipping oligo-1 | CTGTCCATCACGCAAATTAACCGTTGTAGCAATACTTCTTTGATTAGTAATAACATCACCA<br>CGAATTCTCTGCCTCCCTTTTAACCCTAG |
| 48% GC, 1.1 µm loop, unzipping oligo-2 | CTAGGGTTAAAAGGGAGGCAGAGAATTCGTGGTATTAAGAGGCTGAGACTCCTCAAGAGAA<br>GGATTAGGATTAGCGGGGTTTTGCTCAGT |
| 31% GC, 1.1 µm loop, unzipping oligo-1 | TCTGTCCATCACGCAAATTAACCGTTGTAGCAATACTTCTTTGATTAGTAATAACATCACC<br>TCAAATATCAAACCCTCAATCAATATCT |
| 31% GC, 1.1 µm loop, unzipping oligo-2 | AGATATTGATTGAGGGTTTGATATTTGAGGTATTAAGAGGCTGAGACTCCTCAAGAGAAGG<br>ATTAGGATTAGCGGGGTTTTGCTCAGTA |
| 48% GC, 0.65 µm loop, unzipping oligo-1 | TTCGACAACTCGTATTAAATCCTTTGCCCGAACGTTATTAATTTTAAAAGTTTGAGTAACA<br>CGAATTCTCTGCCTCCCTTTTAACCCTAG |
| 48% GC, 0.65 µm loop, unzipping oligo-2 | CTAGGGTTAAAAGGGAGGCAGAGAATTCGTGTCAACCGATTGAGGGAGGGAAGGTAAATAT<br>TGACGGAAATTATTCATTAAAGGTGAATT |
| 31% GC, 0.65 µm loop, unzipping oligo-1 | ATTCGACAACTCGTATTAAATCCTTTGCCCGAACGTTATTAATTTTAAAAGTTTGAGTAAC<br>TCAAATATCAAACCCTCAATCAATATCT |
| 31% GC, 0.65 µm loop, unzipping oligo-2 | AGATATTGATTGAGGGTTTGATATTTGAGTCAACCGATTGAGGGAGGGAAGGTAAATATTG<br>ACGGAAATTATTCATTAAAGGTGAATTA |

**Supplementary Table 1.** Unzipping oligos sequences for DNA construct.
